# Synaptotagmin rings as high sensitivity regulators of synaptic vesicle docking and fusion

**DOI:** 10.1101/2021.03.12.435193

**Authors:** Jie Zhu, Zachary A. McDargh, Feng Li, Shyam Krishnakumar, James E. Rothman, Ben O’Shaughnessy

## Abstract

Synchronous release at neuronal synapses is accomplished by a machinery that senses calcium influx and fuses the synaptic vesicle and plasma membranes to release neurotransmitters. Previous studies suggested the calcium sensor Synaptotagmin (Syt) is a facilitator of vesicle docking and both a facilitator and inhibitor of fusion. On phospholipid monolayers, the Syt C2AB domain spontaneously oligomerized into rings that are disassembled by Ca^2+^, suggesting Syt rings may clamp fusion as membrane-separating “washers” until Ca^2+^-mediated disassembly triggers fusion and release (Wang *et al.*, 2014). Here we combined mathematical modeling with experiment to measure mechanical properties of Syt rings and to test this mechanism. Consistent with experiment, the model quantitatively recapitulates observed Syt ring-induced dome and volcano shapes on phospholipid monolayers, and predicts rings are stabilized by anionic phospholipid bilayers or bulk solution with ATP. The selected ring conformation is highly sensitive to membrane composition and bulk ATP levels, a property that may regulate vesicle docking and fusion in ATP-rich synaptic terminals. We find the Syt molecules hosted by a synaptic vesicle oligomerize into a halo, unbound from the vesicle, but in proximity to sufficiently PIP2-rich plasma membrane (PM) domains the PM-bound trans Syt ring conformation is preferred. Thus, the Syt halo serves as landing gear for spatially directed docking at PIP2-rich sites that define the active zones of exocytotic release, positioning the Syt ring to clamp fusion and await calcium. Our results suggest the Syt ring is both a Ca^2+^-sensitive fusion clamp and a high-fidelity sensor for directed docking.

**Significance:** Synchronous neurotransmitter release relies on directed docking of synaptic vesicles at active zones in axon terminals, where calcium influx activates membrane fusion and release. In vitro, the calcium sensor Synaptotagmin oligomerizes into rings disassembled by calcium. Here, experiment and modeling suggest the Synaptotagmin molecules hosted by an undocked vesicle oligomerize into a tethered, unbound halo in ATP-rich synaptic terminals. The halo directs vesicle docking to PIP2-rich plasma membrane domains in active zones, where the *trans*-bound ring conformation is favored, interposed between the membranes to clamp fusion until calcium triggers ring disassembly and neurotransmitter release. The mechanism exploits the extreme sensitivity of Synaptotagmin ring binding preferences to solution and membrane composition, with ~15 -fold-enhanced sensitivity for rings of ~15 molecules.

## Introduction

Neurotransmission is based on Ca^2+^-triggered release of neurotransmitters from membrane-enclosed synaptic vesicles, each carrying ~ 15 copies of the transmembrane protein synaptotagmin-1 (Syt) (1–3). During synchronous release, an action potential triggers Ca^2+^ entry through voltage-gated Ca^2+^ channels that is sensed by Syt, provoking fusion of the synaptic vesicle with the plasma membrane (PM) and neurotransmitter release. SNARE proteins fuse the vesicle and target membranes, regulated by Syt and other accessory proteins, but the detailed mechanisms remain unclear (4, 5).

The ability of Syt to bind phospholipid membranes in both a Ca^2+^-independent and a Ca^2+^-dependent manner is likely fundamental to its function in synaptic release (6). Syt consists of a transmembrane domain (TMD), a juxtamembrane linker domain (LD) and two Ca^2+^-binding domains, C2A and C2B, that extend away from the vesicle (Fig. 1*A*). The polylysine patch of C2B (residues 324-327) electrostatically binds anionic phospholipids such as phosphatidylserine (PS), with particularly strong binding to phosphatidylinositol 4,5-bisphosphate (PIP2) (6, 7), in a Ca^2+^-independent manner. With Ca^2+^, partial insertion of the calcium-binding loops of the C2 domains into membranes leads to strong Syt-membrane binding (8–10).

**Fig. 1.**
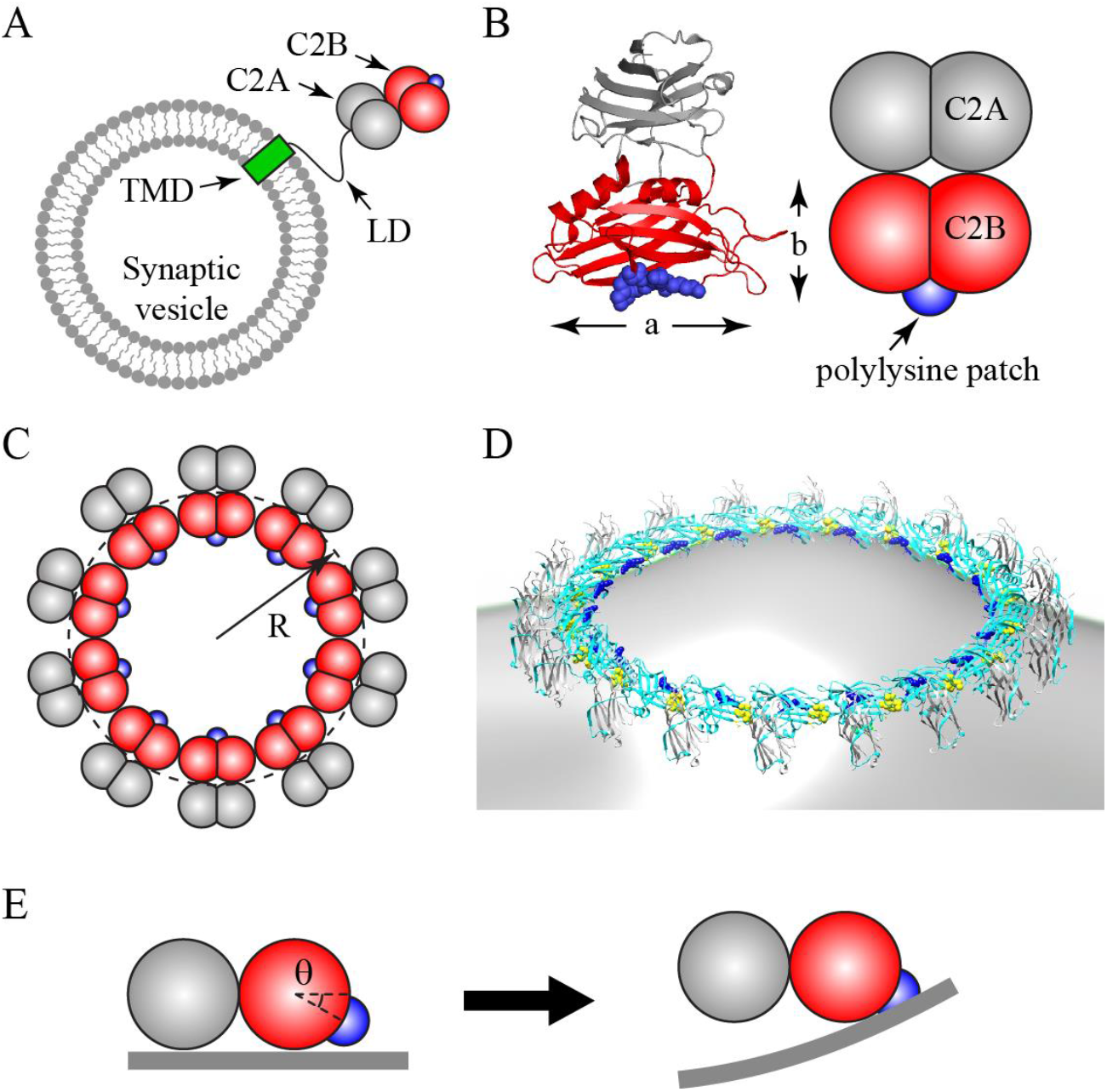
Coarse-grained model of Synaptotagmin (Syt) C2AB domain and oligomeric Syt rings. Synaptic vesicles carry ~ 15 copies of Syt, each comprising a transmembrane domain (TMD), a flexible juxtamembrane linker domain (LD) and the C2A and C2B globular domains (schematic, not to scale). The polylysine patch (blue) on C2B mediates Ca^2+^-independent binding to anionic membranes. (*B*) Crystal structure of Syt (PDB ID: 2R83) (left) and our coarse-grained representation (right). Each C2 domain is represented as two overlapping beads, radius *R*_bead_, giving length *a* = 5 nm and width *b* = 3 nm. The positively charged polylysine patch (blue) on C2B is treated as a point charge. (*C*) Coarse-grained model representation of a Syt ring of radius *R*, top view. (*D*) Rendition of a Syt ring bound to a membrane, based on EM reconstruction of Syt oligomers assembled on phospholipid monolayer tubes (43) (Cyan: C2B. Gray: C2A. Blue: polylysine patch. Yellow: Ca^2+^ binding loops on C2B). (*E*) C2AB subunit in a ring interacting with planar membrane (side view). The polylysine patch lies below the plane of the ring, at an angle *θ* = 30° suggested by EM reconstruction, (*D*). In the ring the C2AB unit cannot rotate downwards, so electrostatic attraction bends the charged membrane up toward the patch (right).

Syt facilitates synchronous fusion, as deletion of Syt shifted neurotransmitter release from a synchronous mode to an asynchronous mode in mice (11, 12) and Drosophila (13). Syt plays a role in clamping fusion before Ca^2+^ entry. Deletion of Syt greatly increased spontaneous release in mice (14, 15) and Drosophila (16–18). The mechanism is unknown, but it was proposed that Syt may clamp fusion in the absence of Ca^2+^ by imposing a membrane separation too great for SNARE-mediated fusion (19), or by binding SNAREs and locking them in a fusion-incompetent state (20–22). Whatever the mechanism, a critical role is likely played by Syt-SNARE binding, recently characterized by structural studies (23–25).

Syt also plays a role in vesicle docking. Syt promoted docking of liposomes *in vitro* (26, 27) and of synaptic vesicles in Drosophila (18, 28), in mouse hippocampal neurons (29, 30), and in mouse chromaffin cells (31). Ca^2+^-dependent *trans*-binding of Syt to PIP2-containing liposomes from Syt-bearing chromaffin granules was reported, as well as *cis*-binding to the host granules (32, 33). Importantly, the presence of ATP reduced the *cis*-binding but not the *trans*-binding (32). Synaptotagmin mutants lacking the polylysine patch docked vesicles deficiently to the PM before arrival of an action potential (29).

Vesicle fusion occurs at active zones in the presynaptic membrane with high PIP2 content (19, 34). RIM proteins and Munc13 are known to play essential roles in the localization of synaptic vesicles to active zones (35, 36), but the detailed mechanisms that target vesicles to these high PIP2 regions remain uncertain (31, 36, 37). Rapid vesicle fusion requires PIP2 (38, 39). PIP2 regulates fusion machinery components such as Syt to be more effective as facilitators of fast fusion and release (40). The speed at which Syt responds to calcium by penetrating the membrane is enhanced by the presence of PIP2 (41), and PIP2 enhances the sensitivity of Syt to calcium 40-fold (7, 42).

In the absence of Ca^2+^, Syt oligomerizes into ring-like oligomers. Without ATP, electron microscopy (EM) images showed rings of Syt^C2AB^, the soluble C2AB domain, on carbon-supported anionic phospholipid monolayers (43, 44) that buckled into dome-, volcano- or tube-like shapes. Ring formation depended on the polylysine patch of the C2B domain, as the KAKA (K326A, K327A) patch mutation abolished rings (43, 44). Higher PS or PIP2 membrane content or lower salt concentration increased the ring density.

These findings suggest Syt^C2AB^ ring formation is mediated by electrostatic interactions between anionic lipids and the polylysine patch of C2B. Rings appear further promoted by the highly charged LD (45), since Syt^CD^, the cytosolic domain including the LD, oligomerized into rings on monolayers even at physiological salt concentrations and in the presence of ATP, while Syt^C2AB^ rings required low salt (44). In the presence of ATP or soluble PIP2, Syt^C2AB^ rings can also assemble in solution, at physiological and lower salt concentrations (46).

Importantly, membrane-bound Syt rings disassemble at physiological Ca^2+^ concentrations (43), suggesting an explanation for Syt’s dual role as fusion clamp and facilitator. In this model, SNAREs are primed for fusion prior to Ca^2+^ influx, but the Syt ring acts as a “washer” separating the synaptic vesicle and plasma membranes, sterically preventing fusion; injection of Ca^2+^ rapidly disassembles the ring and fusion proceeds (43). Consistent with a clamping role for Syt oligomers, mutations selectively disrupting Syt oligomerization show increased spontaneous release frequency in PC12 cells (47) and neocortical synapses (48) and abrogated clamping under *in vitro* conditions (49, 50). This mutation also abolished a symmetric arrangement of components revealed by electron cryotomography at the synaptic vesicle-plasma membrane interface in nerve growth factor-differentiated PC12 cells, suggesting Syt rings template the organization and have a role consistent with this model (51). Ca^2+^ binding may disassemble rings by driving Ca^2+^-dependent insertion of the Ca^2+^-binding loops into membranes, as Ca-triggered disassembly is abolished by the 3xDA (D309A, D363A, D365A) mutation, which disrupts Ca^2+^ binding to the C2B domain (43, 44).

Here we assess the feasibility of the hypothesized Syt washer mechanism quantitatively, combining molecularly explicit mathematical modeling and experiment. We measured Syt ring size distributions in bulk solution, and used a mathematical model to infer the bending stiffness of rings. We built a model of Syt-Syt and Syt-membrane interactions severely constrained by experimental data, which quantitatively reproduced the novel Syt-induced monolayer domes and volcanoes. Applying the model to the conditions in synaptic terminals during neurotransmitter release when vesicles hosting ~ 15 copies of Syt dock at the PM, we asked if the Syt C2 domains will oligomerize into a ring and, if so, whether the ring will bind the PM as assumed by the washer model (*trans*-binding), bind the vesicle membrane (*cis*-binding) or remain in solution (the “halo”). We find the outcome depends with extreme sensitivity on membrane and cytosol composition. These results suggest the Syt halo serves as landing gear for spatially directed vesicle docking to PIP2-rich sites that colocalize with the t-SNARE Syntaxin and other fusion machinery components in active zones, positioning the PM-bound trans Syt ring to clamp fusion and await calcium. Thus, the Syt ring is both a Ca-sensitive fusion clamp and a high-fidelity sensor for directed docking. The high sensitivity originates in the oligomeric character of the ring, which enhances sensitivities ~ 15-fold compared to monomeric Syt.

## Results

### Model of Syt oligomerization in solution to extract mechanical properties of Syt rings

We used electron microscopy (EM) to image Syt C2AB domains incubated in solution with 15 mM KCl and 1 mM ATP (see *Materials and Methods*). To ease notation, hereafter Syt will denote the C2AB domain (referred to as Syt^C2AB^ above). Negative stain analysis showed that Syt monomers in solution spontaneously oligomerized into rings. We measured the size distribution, revealing a mean radius *R* = 13.1 ± 1.4 nm (mean ± s.d.) (Fig. 2*A*).

**Fig. 2.**
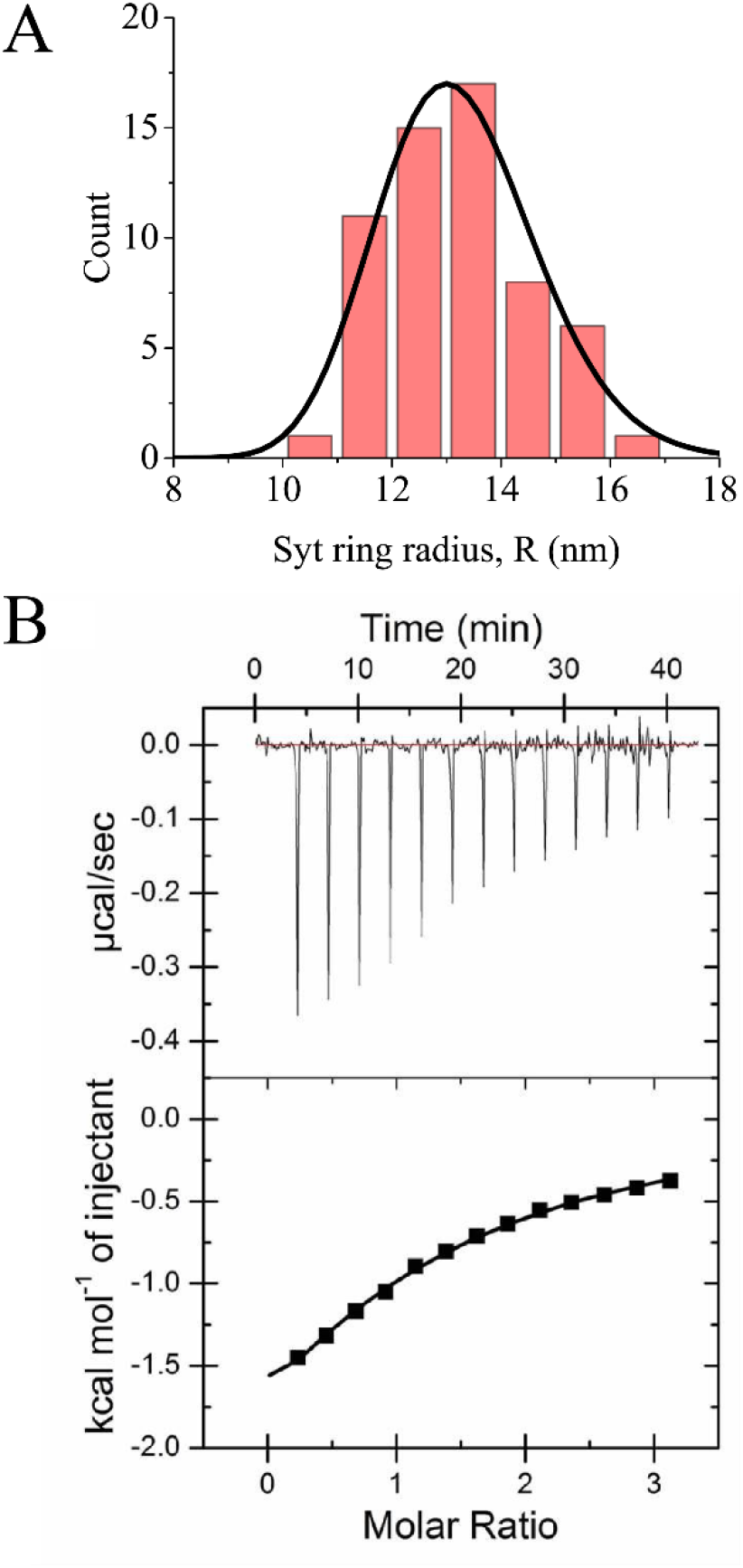
(*A*) Experimental Syt ring size distribution in bulk solution with 1mM Mg-ATP and 15 mM KCl, from electron micrographs (radius *R* is defined in Fig. 1*C*). Mean radius is 13.1 nm ± 1.4 nm (S.D.). Solid line: best fit theoretical distribution (Eq. 1), yielding spontaneous radius of curvature *R*_0_ = 13 nm and persistence length *l*_*p*_ = 170 nm (Table 1). *(B)* Syt-ATP binding measured by Isothermal Titration Calorimetry (ITC). The measured parameters are: H = −4.4 ± 0.7 kcal/mol, dissociation constant 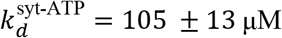.

To extract the stifness and energetics of Syt rings from this size distribution, needed to model syt ring formation and interaction with membranes (next section), we developed a simple model describing the equilibrium of monomers and rings in solution. Rings were assumed circular, with variable numbers of Syt monomers each of length *a* = 5 nm, spontaneous radius of curvature *R*_0_ and unknown flexural rigidity *l*_*p*_kT, where *l*_*p*_ is the persistence length (52). We find the effective free energy of rings of radius *R* is given by (*Supporting Information*),

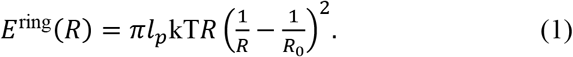

This expression involves neither the Syt-Syt binding energy *∈*^syt−syt^ nor the Syt monomer concentration, since in equilibrium the latter self-tunes to a critical value where the free monomer chemical potential equals *∈*^syt−syt^ (*Supporting Information*). The predicted ring size distribution is the Boltzmann distribution, *P*^ring^(*R*)~exp[−*E*^ring^(*R*)/kT] (solid line, Fig. 2*A*). Fitting to the experimental distribution yielded best fit values *R*_0_ = 13 nm, *l*_*p*_ = 170 nm, or ~16 Syt subunits per ring, in agreement with fluorescence correlation spectroscopy measurements (46) and similar to the number of Syt molecules per synaptic vesicle, ~15-20 (1, 3). Overall, we find Syt rings are quite flexible; e.g., a ring of 20 subunits has only 1.7 kT more free energy than a ring with the mean number, 16.

### Coarse-grained model of Syt rings interacting with membranes

To describe the Ca^2+^-independent interaction of Syt rings with anionic phospholipid membranes, we developed a highly coarse-grained model, as system size and equilibration timescales are considerable. The main model outputs are: (i) the membrane surface configuration with a bound ring, and (ii) the free energy of ring-membrane binding. Here we present the framework, which will be adapted to monolayers and then to physiologically realistic bilayers in later sections (see *Supporting Information*).

A Syt^C2AB^ molecule comprises a C2A and a C2B domain. The crystal structure of human Syt (PDB ID: 2R83) shows the domains are similarly shaped, about *a* = 5 nm long and *b* = 3 nm wide (Fig. 1*B*). We represent each C2 domain by two beads of radius *R*_bead_ = 1.5 nm, overlapping to give length *a* and width *b* (Fig. 1*B* and *Supporting Information*).

Syt rings are constructed by interconnecting many Syt^C2AB^ subunits. From EM reconstruction of Syt oligomers on monolayer tubes (43), the C2B domains form the inner circle of the ring, their long axes oriented tangentially (Figs. 1*C*,*D*). The C2B polylysine patches lie on the inward face of the ring, but tilted an angle *θ* = 30° out of plane (Fig. 1*E*). Thus, critically, when a ring binds a membrane, the membrane must curve upwards to bind this patch.

Phospholipid membranes were modeled as continuous sheets using a well-established triangulation scheme with a dynamic mesh representing membrane fluidity, bending energy and tension (53–55) (see *Supporting Information*). The membrane sheet is a network of nodes, with adjacent nodes connected by finitely extensible tethers and interacting via hard-sphere repulsions below a certain separation. Random bond flipping ensures fluidity, continuously redefining node-node connectivity. Bending and tension are discretized versions of the Helfrich energy (56),

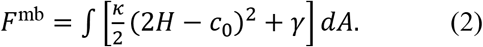

Here *k* is the bending modulus, *H* the mean of the principal curvatures, discretized using the scheme of ref. (57), *c*_0_ is the spontaneous curvature, and *γ* the membrane tension.

Syt interacts with the membrane via screened electrostatic and repulsive steric forces. Membranes are assumed uniformly charged, so each node interacts electrostatically with the polylysine patch of each Syt with a strength depending on membrane composition. Taking the reference state as free Syt monomers in solution, the total free energy of formation of a Syt ring bound to a membrane, per Syt monomer, is given by

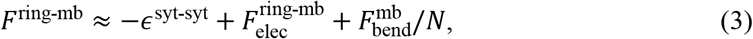

where *∈*^syt−mb^ denotes the Syt binding energy, 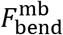 the bending energy and 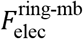 the bend elec electrostatic interaction between the Syt ring and the membrane, all per monomer. The superscript “mb” may denote “pm”, “ves” or “mono” depending on whether the planar or vesicle membrane or monolayer is involved. In Eq. (4), we neglected contributions from membrane tension, Syt-membrane steric interactions and Syt ring bending energy (Eq. 1), all of which our simulations showed to be negligible.

To equilibrate, we placed a Syt ring adjacent to a flat 60 nm diameter monolayer or 50 nm diameter bilayer, and evolved the system using pseudo Langevin dynamics (58) which decrease free energy with time (*Supporting Information*) using interaction forces from energy derivatives and fictitious particle drags. In simulations with vesicles, the 40 nm diameter vesicles had pressure-enforced constant volume.

### Measurement or calculation of Synaptotagmin binding parameters needed by the model

We first experimentally measured or computed key model parameters (see Table 1 and *Supporting Information* for details).

**Table 1.** Model parameters.

| Symbol |  | Value | Source |
| --- | --- | --- | --- |
| $a, b$ | Length, width of C2B domain | 5 nm, 3 nm | (43) |
| $c_0$ | Monolayer spontaneous curvature (40% PS, 60% PC) | $0.02 \text{ nm}^{-1}$ | (A) |
| $d_h$ | Hydrophobic interaction length | 2 nm | (59, 60) |
| $E_h^0$ | Monolayer-carbon support hydrophobic energy density | $0.04 \text{ kT/nm}^2$ | (B) |
| $\epsilon^{\text{syt-ATP}}$ | Syt-ATP binding energy | 9.7 kT | (C) |
| $\epsilon^{\text{syt-mono}}$ | Syt-monolayer binding energy | 18 kT | (D) |
| $\epsilon^{\text{syt-pm}}$ | Syt-planar-membrane binding energy | 4.9 kT | (D) |
| $\epsilon^{\text{syt-syt}}$ | Syt-Syt binding energy | 9.5 kT | (E) |
| $\epsilon^{\text{syt-ves}}$ | Syt-synaptic-vesicle binding energy | 2.2 kT | (D) |
| $l_p$ | Persistence length of Syt oligomer | 170 nm | (F) |
| $R_{\text{bead}}$ | Radius of model beads | 1.5 nm | (G) |
| $R_{\text{ves}}$ | Vesicle radius | 20 nm | |
| $R_0$ | Spontaneous radius of curvature of Syt oligomers | 13 nm | (F) |
| $\kappa$ | Bending modulus of monolayer (bilayer) | 12.5 kT (25 kT) | (H) |
| $\gamma$ | Bilayer membrane tension | 0.003 pN/nm | (75, 76) |
| $\eta$ | Viscosity of water | $10^{-3} \text{ Pa}\cdot\text{s}$ | |
| $\eta_m$ | 2D viscosity of lipid bilayers | $3 \times 10^{-9} \text{ Pa}\cdot\text{s}\cdot\text{m}$ | (77) |
| $\lambda$ | Debye length at 140 mM salt | 0.7 nm | (65) |

**Notes:** (A) Linear interpolation of spontaneous curvatures of DOPS ( $0.07 \text{ nm}^{-1}$ (78)) and DOPC ( $-0.02 \text{ nm}^{-1}$ (79)), based on the molar fractions of the two components.
(B) Deduced in the present work (see Figs. 3C,D).
(C) Experimentally measured in the present work.
(D) Calculated in the present work, using measurements of ref. (6) as input (see main text).
(E) Calculated in the present work, using measurements of ref. (46) as input (see main text).
(F) Calculated in the present work by fitting Eq. 1 to the experimentally measured Syt ring size distribution in bulk solution (Fig. 2A).
(G) Estimated from the C2AB crystal structure (Fig. 1B).
(H) Bilayer bending modulus from ref. (80), measured in 100 mM NaCl. Monolayer modulus is assumed half the bilayer value.

**(1) Syt-ATP binding energy, Fig. 2*B***. We used isothermal titration calorimetry (ITC) to measure the dissociation constant 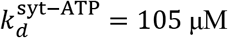 at 140 mM KCl. Taking an ATP molecule size *a*~1 nm gives a local binding energy 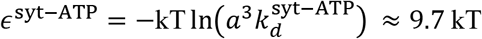. **(2) Syt-Syt binding energy in the presence of 1 mM Mg-ATP.** Wang et al. measured 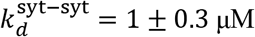 with 20 mM KCl and 1 mM Mg-ATP, yielding binding energy *∈*^syt−syt^ ≈ 9.5 kT, taking a Syt molecule size 5 nm (46). We assumed this value is independent of salt concentration and unaltered by membrane binding. **(3) Syt binding to phospholipid bilayers and monolayers in the absence of Ca^2+^.** Using the molar partition coefficient *K* = 160 M^−1^ reported for Syt-liposome binding (25% POPS, 75% POPC) in 100 mM KCl (6), we extrapolated the binding energy to different salt concentrations and membrane charge densities. For physiological conditions this yielded ∈^syt−pm^ ≈ 4.9 kT and *∈*^syt−ves^ ≈ 2.2 kT for a typical neuronal PM and synaptic vesicle, respectively, and *∈*^syt−mono^ ≈ 18 kT for the monolayers studied here (43) (see Table 2 for assumed membrane compositions).

**Table 2.**
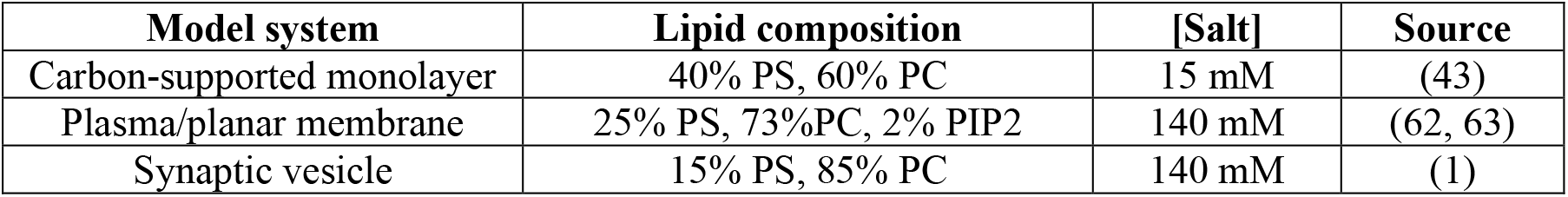
Lipid composition and salt concentration conditions used by model

| **Model system** | **Lipid composition** | **[Salt]** | **Source** |
| --- | --- | --- | --- |
| Carbon-supported monolayer | 40% PS, 60% PC | 15 mM | (43) |
| Plasma/planar membrane | 25% PS, 73%PC, 2% PIP2 | 140 mM | (62, 63) |
| Synaptic vesicle | 15% PS, 85% PC | 140 mM | (1) |

### Syt oligomerizes into rings on phospholipid monolayers and generates domes or volcanoes

We first asked if our model could reproduce the experiments of (43), in which Syt rings formed on carbon-supported monolayers and the monolayer assumed various shapes at the ring locations. To mimic experiment, we simulated monolayers at 15 mM KCl (Table 1) and with uniform charge density corresponding to the lipid composition of 40% PS, 60% PC (Table 2). Free energies were calculated from Eq. (4), but with an additional term added to *F*^ring-mb^ of Eq. (4) representing the hydrophobic attraction (per Syt monomer) between the monolayer and its carbon support (59, 60), 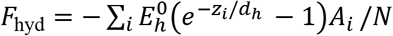. Here 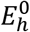 is the hydrophobic energy density, *Z*_i_ the separation ℎ ℎ between the *i*^th^ monolayer node and the carbon substrate, *A*_*i*_ the area of node *i*, and *d*_*h*_ ≈ 2 nm the hydrophobic decay length.

Placing the ring adjacent to the monolayer as initial condition, independently of initial condition the system evolved to an equilibrium with the Syt ring bound to the monolayer which it deformed into a dome or volcano, similar to experiment (43) (Fig. 3*A*,*B*). The selected shape depended on the number of Syt molecules per ring, *N*, and the hydrophobic interaction 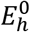 (see phase diagrams, Figs. 3*C*,*D*)). Volcanoes are favored by large rings and strong hydrophobic attraction.

**Fig. 3.**
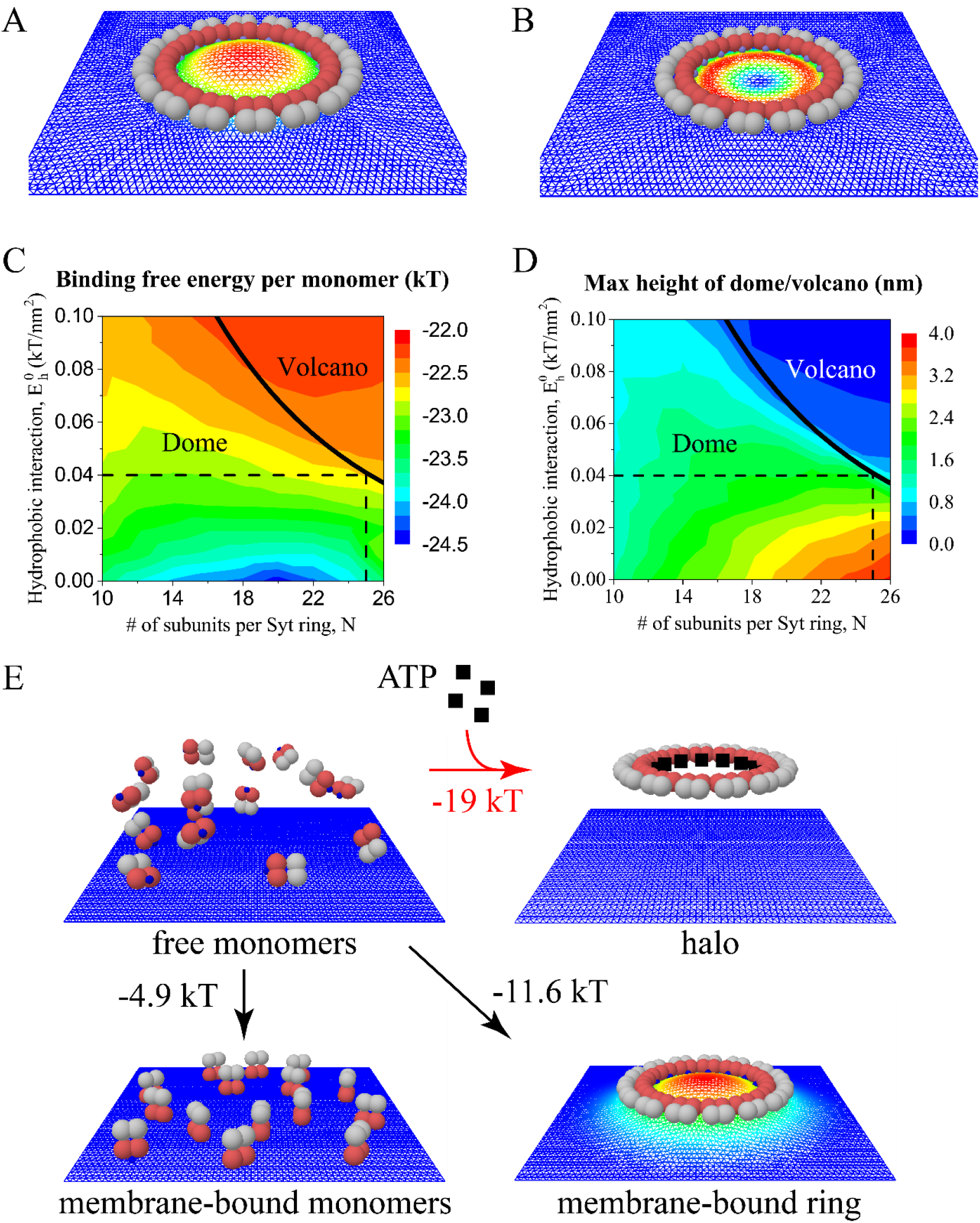
Model results for binding of Syt rings on carbon-supported monolayers and on planar bilayer membranes. (*A*-*B*) Simulation snapshots of equilibrium configurations of a Syt ring of *N* = 16 subunits binding a monolayer with 40% PS, 60% PC in 15 mM salt solution (conditions as in ref. (43)). Warmer colors denote greater monolayer elevations. (*A*) Monolayer deforms into a dome shape for lower monolayer-carbon hydrophobic interaction energy, 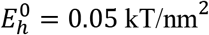. Volcano shape results for larger interaction, 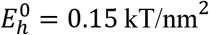. (*C-D*) Monolayer shape (dome or volcano) and binding free energy per monomer, (*C*), or maximum monolayer height, (*D*), versus *N,* the number of Syt subunits per ring, and 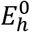, the monolayer-carbon binding energy density. In the experiments of ref. (43), the dome-volcano transition occurred at *N* ≈ 25, identifying 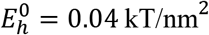 (dashed lines, see main text). (*E*) Possible states for Syt monomers in solution near a planar bilayer membrane with overall composition typical of a plasma membrane, composition assumed homogenous (Table 2). Energies of formation are per Syt monomer, relative to a reference state with 15 free Syt monomers in solution at physiological salt conditions, 140 mM (top left). With ATP, the preferred state is the Syt ring in solution (halo), stabilized by the substantial Syt-ATP binding energy, *∈*^syt−ATP^ ≈ 9.7 *kT*. Without ATP the Syt ring oligomerizes on the membrane, deforming it into a dome (simulation snapshot, warmer colors denote greater elevation).

To estimate the hydrophobic interaction strength 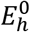, we used the fact that the dome-volcano transition occurred at *N* ≈ 25 in the experiments of Ref. (43), corresponding to 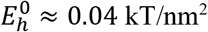on the predicted dome-volcano boundary curve (Fig. 3*C*). With this value, over a range of *N* values the free energy of ring formation is ~ − 25 kT/monomer, with dome and volcano heights ~ 1-2 nm and <1 nm, respectively. The negative free energy shows that Syt spontaneously forms rings on carbon-supported monolayers from solution.

The origin of these shapes is clear from the model. When Syt assembles into a ring, the polylysine patches are raised ~1 nm above the membrane (Fig. 1*C,E*), so that binding requires the membrane to deform upwards everywhere around the ring into a dome shape (Fig. 1*D,E*). This shape incurs a membrane bending energy penalty proportional to ring length, ~*N*, being dominated by the severe curvature at the ring boundary. Another candidate shape, the volcano, has a higher bending energy ~*N*, as the high curvature occurs along both the inside and outside of the volcano rim. However, a dome pulls the monolayer away from the carbon substrate, incurring a hydrophobic penalty ~*N*^2^ proportional to ring area, whereas the hydrophobic penalty is only ~*N* for the volcano. Thus, for large enough *N* the net dome energy exceeds that of the volcano, and volcanoes are selected. The greater 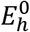, the smaller the value of *N* at the transition. These features are apparent in Fig. 3*C*.

### Syt forms rings on homogenous planar bilayers but rings prefer ATP-rich solution

Our model predicts that Syt C2AB molecules in solution spontaneously oligomerize into monolayer-bound rings, as seen experimentally. We next asked if the same is true for physiologically relevant bilayers, untested experimentally. This is a priori unclear, as the hydrophobic substrate interaction is absent (favoring ring binding), but a bilayer has twice a monolayer’s bending modulus (disfavoring ring binding). Further, the ~1 mM ATP in the cytosol (61) may favor rings in solution an effect seen for carbon-supported monolayers with physiological ATP concentrations (44, 46). All energies below are per Syt monomer, unless otherwise stated.

We simulated Syt rings interacting at physiological salt concentration with planar bilayers of typical PM composition (Table 1), including 25% PS and 2% PIP2 (62, 63) (Table 2). The composition was assumed homogenous. Rings of 10-20 subunits bound to the bilayer, producing domes (Fig. 3*E*). In the final state the attractive electrostatic energy ~2.5 kT was offset by a membrane bending energy penalty of ~0.4 kT. Adding the Syt-Syt binding energy ~9.5 kT, we conclude that Syt monomers in ATP-free solution bind bilayers as rings, with net binding free energy *F*^ring-pm^ ≈ −11.6 kT per monomer. This has greater magnitude than the binding energy of free monomers from solution, *∈*^syt-pm^ ≈ 4.9 kT. Thus, membrane-bound rings are stable to fragmentation into membrane-bound monomers (Fig. 3*E*).

Consider now the effects of ATP in the bulk solution. Thus far we excluded Syt rings in solution (“halos”) as candidate structures. With ATP we find halos are preferred, since Syt molecules can release *∈*^syt−ATP^ ≈ 9.7 *kT* by remaining in solution and binding ATP. Thus, the energy of halo formation is *F*^halo^ = −(*∈*^syt-ATP^ + *∈*^syt-syt^) ≈ −19kT, less than that of a membrane-bound ring, *F*^ring-pm^ (Fig. 3*E*). This argument assumes Syt in solution must unbind ATP in order to bind the membrane (32, 44).

### Membrane curvature and charge density effects: Syt forms rings that bind anionic vesicles in the absence of ATP

The previous section treated Syt interactions with a model PM. At synaptic terminals, Syt can presumably also interact with its high curvature host synaptic vesicle, lacking PIP2 (1) (Table 2). Thus, we analyzed curvature and charge density effects.

Simulations showed that Syt monomers oligomerize into rings and bind anionic vesicles when ATP is absent. At physiological salt, Syt rings (10 ≤ *N* ≤ 20) bound and dimpled 20 nm radius vesicles with synaptic vesicle lipid composition, a milder version of the dome on planar membranes (Fig. 4*A*). Typical energies were ~2 *kT* (electrostatic advantage) and ~0.2 kT (membrane bending penalty), giving a net binding energy *F*^ring-ves^ ≈ −11.3 kT, after adding the Syt-Syt binding advantage (Fig. 4*B*-*D*).

**Fig. 4.**
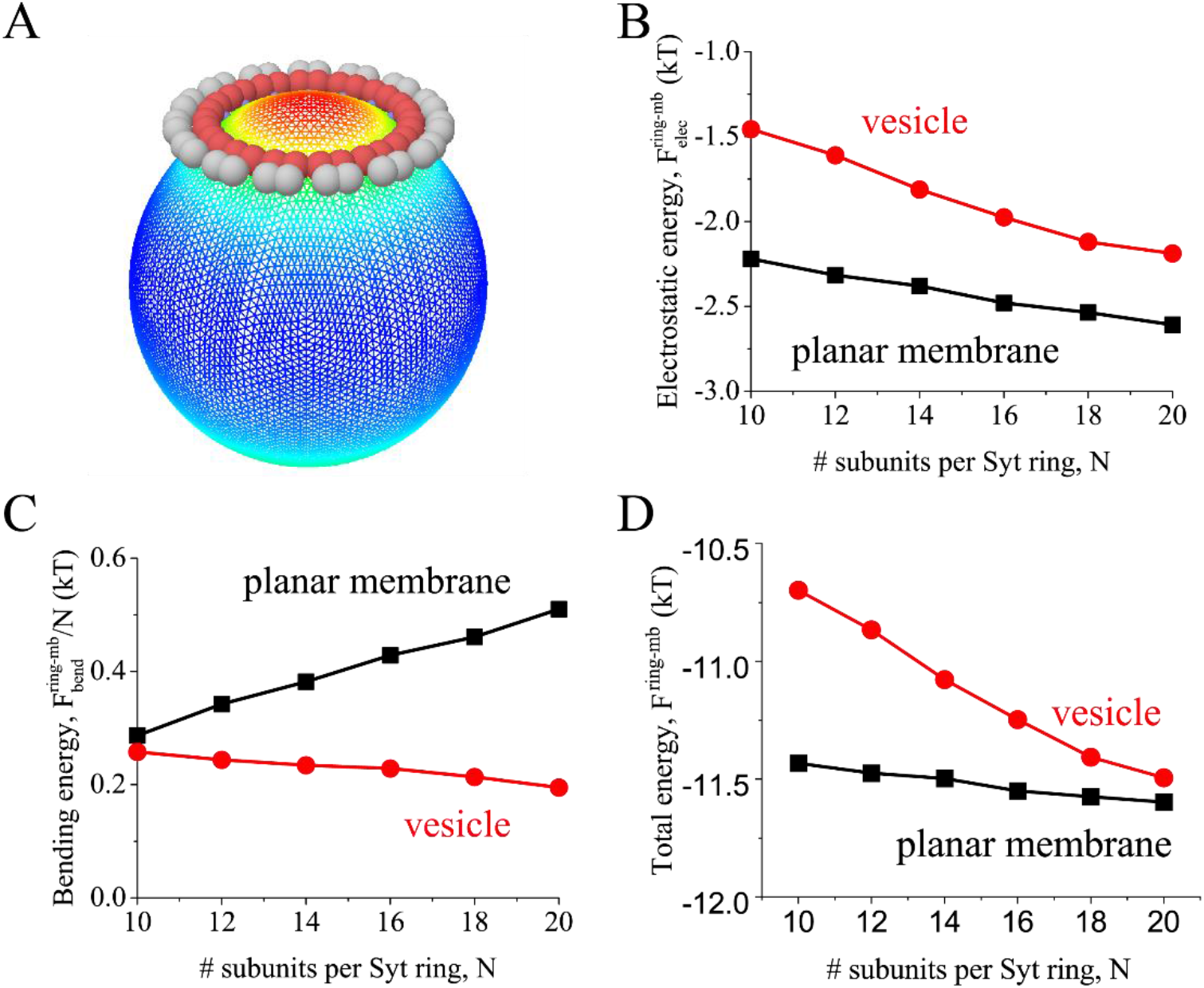
Membrane curvature and charge density promote binding of Syt rings. Model results for binding of Syt rings at physiological salt, 140 mM, to a 20 nm radius lipid vesicle with composition representative of synaptic vesicles (15% PS, Table 2). Vesicle binding energies are compared with binding energies to planar membranes with lipid composition representative of a plasma membrane (Table 2). Energies are per Syt subunit. (*A*) Simulation snapshot of a Syt ring of *N* = 15 subunits binding and dimpling the vesicle, a mild version of the dome seen on planar bilayers. Warmer colors denote greater distance from vesicle center. (*B*-*D*) Model-predicted free energies when a Syt ring binds a vesicle or a planar membrane versus number of Syt monomers per ring. (*B*) The attractive electrostatic energy is greater for the plasma membrane due to its higher anionic lipid charge density. (*C*) Vesicle binding incurs less bending energy, due to vesicle curvature. (*D*) The net binding energy favors the plasma membrane, since the electrostatic advantage exceeds the bending disadvantage.

The competition between a planar PM and a vesicle for binding to a Syt ring is quantified in Fig. 4*B-D*. Vesicle curvature favors ring binding, because the membrane needs to bend less to curve upwards and bind the polylysine patch (Fig. 1*E*). However, electrostatic interactions favor Syt-PM binding due to the higher PM negative charge density. Overall, binding to planar PMs is preferred (Fig. 4*D*). We stress, however, that the planar membrane energies of Fig. 4 neglect composition inhomogeneities, which are considered next.

### PIP2 clustering promotes *trans*-binding of Syt rings to the plasma membrane

In synaptic terminals, ~15 Syt^C2AB^ monomers are anchored to a synaptic vesicle by their TMDs and juxtamembrane LDs (1) and ~1 mM ATP (61) is present. When a vesicle approaches the PM in an active zone, if the monomers oligomerize into a ring, the ring could bind the PM (*trans*-binding) as predicted by the washer hypothesis, bind the vesicle membrane (*cis*-binding) or exist unbound in solution (halo) (Fig. 5*A*).

**Fig. 5.**
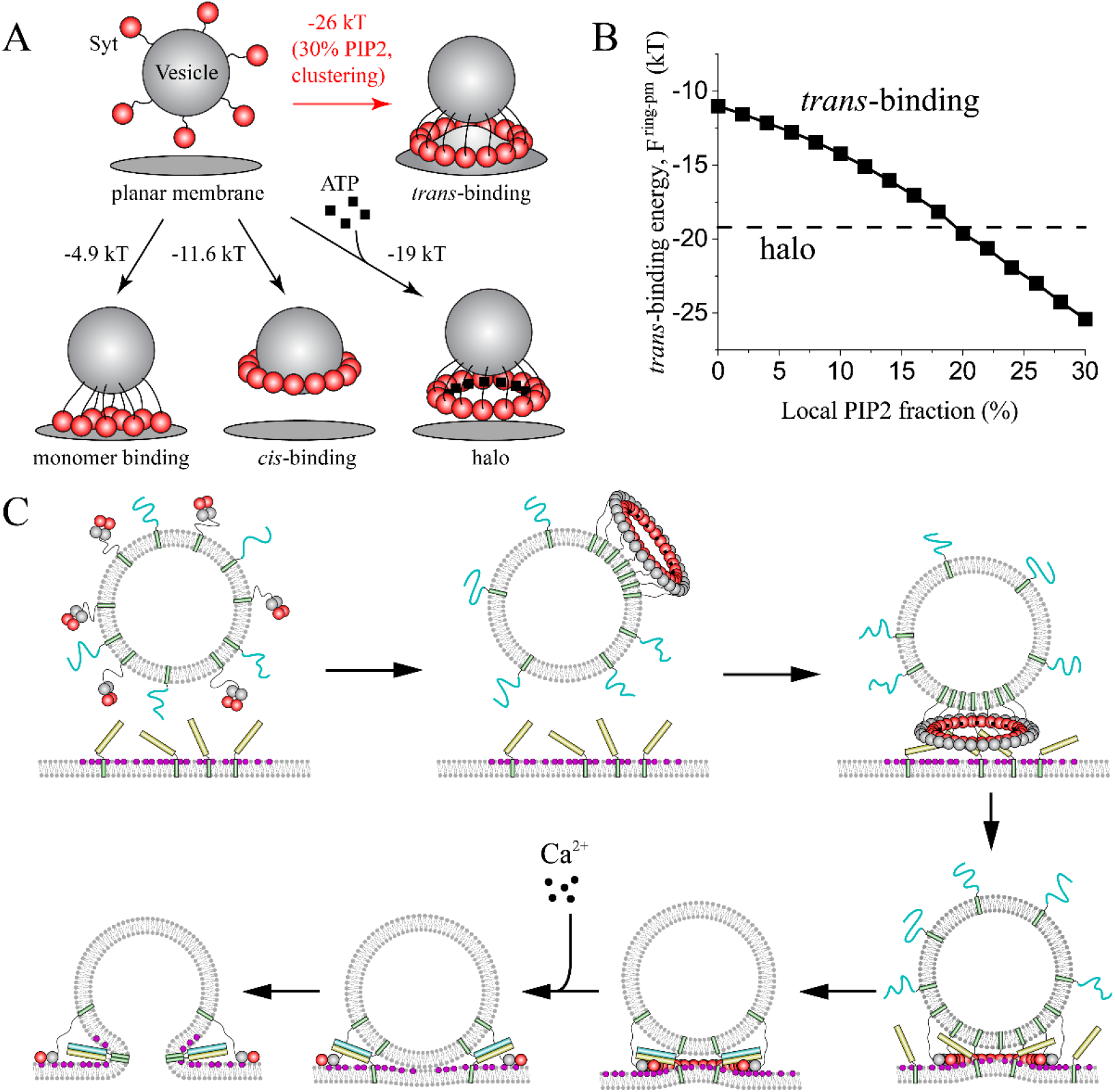
(*A*) When a synaptic vesicle bearing 15 Syt molecules approaches the plasma membrane, four final states are possible. The model-predicted free energies per molecule are shown. Vesicle size, lipid composition and salt conditions, as for Fig. 4. At PIP2 clustering sites with sufficiently high local PIP2 concentration *trans*-binding of a Syt ring is preferred (e.g. 30% PIP2, as shown) to the ATP-stabilized halo, the *cis*-bound ring or unoligomerized membrane-bound monomers. (*B*) Model calculations of *trans*-binding energy of a Syt ring to the plasma membrane versus local PIP2 concentration. For [PIP2] > 18%, *trans*-binding is favored over the halo, while halos are preferred for [PIP2] < 18%. (*C*) Model of Syt-mediated regulation of neurotransmitter release, with the Syt ring serving both as a discriminating sensor and as a spacer clamping fusion. In the ATP-rich synaptic terminal, the ~15 Syt^C2AB^monomers carried by a synaptic vesicle form a halo (unbound ring) in preference to a ring bound to the PIP2-free vesicle (*cis*-binding). The oligomeric nature of the Syt ring endows it with high sensitivity to PIP2 levels in the plasma membrane, and trans-binding to the PM is preferred to the halo only for PIP2 content above a threshold. Thus, the Syt halo is spatially directed to dock the vesicle at PIP2-rich sites which colocalize with the t-SNARE Syntaxin (yellow). Docking positions the Syt ring to clamp fusion by spacing the vesicle and PMs, and by restraining the SNARE proteins to lie partially within the Syt ring, consistent with measured crystal structures of the Syt-SNARE complex (23, 24). Ca^2+^ influx disassembles the ring, releasing SNAREs to fuse the membranes rapidly and release neurotransmitters through the fusion pore.

Our results so far show that Syt rings bind the PM without ATP, but form halos with ATP, suggesting Syt halos are favored in ATP-rich synaptic terminals. This would seem to argue against the washer hypothesis. However, all calculations thus far assumed uniformly distributed membrane charge, neglecting the established clustering of the highly charged lipid PIP2 (a charge −4e is reported at PH 7 (62, 64)). In the PM, PIP2 clusters into ~70 nm diameter microdomains (65) with local PIP2 concentrations 20-100% (63). Further, the t-SNARE Syntaxin is thought to colocalize with PIP2 in active zones (66), suggesting that vesicle docking and synaptic release occur within these microdomains.

To account for PIP2 inhomogeneities, we next ran simulations with the PM carrying a charge density based on the local (rather than global) PIP2 concentration, modeling a vesicle near a PIP2 “hot spot” in the active zone. In our model, this is achieved by accordingly increasing the Syt-membrane binding energy parameter in the electrostatic energy, *∈*^syt-pm^~[PIP2] (see *Supporting Information*).

Since the precise value of the local [PIP2] is unknown, we ran simulations for a range of values. The ring-PM binding energy increased with increasing local [PIP2] (Fig. 5*B*). For *N* = 15, the *trans*-binding energy of a ring became comparable to the halo energy at [PIP2]≈ 20%. Given the reported local PIP2 fractions of ≥ 20% (63), we conclude that, consistent with the washer hypothesis, *trans*-binding of Syt rings is favored when vesicles approach PIP2-rich PM domains in synaptic terminals. By contrast, we predict that the halo is favored outside PIP2-rich domains. These conclusions are robust with respect to the presence of cholesterol, which considerably increases membrane bending modulus (67), since simulated *trans*-binding energies were increased only ~ 0.3 kT by a 4-fold increase of bending modulus to 100 kT (Fig. S1).

The principal conclusions of this study are summarized in Fig. 5*A*. In ATP-rich synaptic terminals away from PIP2 microdomains halos are favored, having free energy ~ −19 kT. However, near a cluster with even a relatively low [PIP2] concentration of 30%, trans-binding is favored as the trans-binding energy is ~ 7 kT lower than the free energy of halo formation.

## Discussion

### Due to their oligomeric nature Syt rings bind membranes with high sensitivity to lipid composition and membrane curvature

Here we predict that Syt rings prefer to bind membranes with higher anionic lipid density or curvature (Fig. 4), and the preference is characterized by high sensitivity. Thus, rings bind vesicles in preference to planar membranes of the same lipid composition, but planar membranes of sufficiently high charge density are preferred, as at neuronal synapses. The transition in preference is predicted to be very sharp, i.e. Syt rings bind specific targets with high sensitivity, an effect well known from the surface adsorption energetics of high molecular weight synthetic polymers (68). The origin is the *N*-fold greater binding energy of an oligomer of *N* units, compared to one unit. Since the binding energy is compared to a fixed scale ~ *kT*, this translates to *N*-fold higher sensitivity. It would be interesting to test these and other specific predictions experimentally. *In vitro* these predictions might be tested by measurements of interactions of Syt-reconstituted vesicles with suspended bilayers or other vesicles, as a function of vesicle sizes and compositions.

### Ca2+-independent binding of Syt rings to membranes is coupled to membrane bending

Dome or volcano shapes were observed in electron micrographs at sites of Syt ring binding on phospholipid monolayers (43). The model developed here explained these as due to electrostatic attraction of the monolayer to the charged polylysine patches of the C2B domains lining the inner edge of the ring that overcomes the cost of membrane bending energy and bulges the monolayer upwards. Thus, binding is coupled to monolayer bending. We find the same qualitative effect occurs on bilayers (Figs. 3, 4). This mechanism is distinct from a previously proposed Ca^2+^-dependent Syt-mediated membrane bending mechanism, which holds that insertion of the C2B loop leads to local bending (69).

In this study the significance of the domes or volcano shapes was as a quantitative test of the model. Whether they have functional significance is unknown. A dome could bring the membrane into closer proximity to the Syt calcium binding loops to accelerate Syt ring disassembly on injection of Ca^2+^. However, membrane buckling was not resolved beneath docked vesicles in cryo-electron tomographic analysis of PC12 cells and cultured neurons (51, 70), or on supported monolayers when the full cytosolic domain of Syt was included (44).

### Syt halo formation may prepare Syt for its role as a fusion clamp and direct synaptic vesicle docking

ATP promotes Syt ring formation in bulk solution (44). The model presented here predicts that, due to the presence of ATP at synaptic terminals, the ~15 Syt monomers carried by a synaptic vesicle form a tethered halo in preference to a vesicle-bound ring (*cis*-binding). The halo ring is assembled away from the vesicle surface, so that halo formation may prepare the ring for its subsequent role as a *trans*-binding fusion clamp. In this picture the Syt ring is the landing gear for docking, ensuring that once the vesicle is docked the Syt ring is ready-formed and correctly positioned at the fusion site to serve as a spacer and fusion clamp.

Furthermore, we predict that the halo fails to bind phospholipid membranes whose PIP2 content is low, because the halo is then energetically favored compared to a Syt ring bound in *trans* to the PM (Fig. 5). By contrast, in the vicinity of PIP2 clusters with local PIP2 concentrations exceeding a threshold ~20%, *trans*-binding is preferred. (This threshold is likely underestimated, as divalent cations such as Mg^2+^ (64) or highly basic protein residues (63, 66) may partially neutralize clustered PIP2; however, local PIP2 concentrations of up to ~80% have been measured in clusters (63).) Thus, the halo landing gear specifically directs docking to PIP2-rich sites.

These results suggest Syt ring formation may underlie a mechanism to direct the docking of vesicles to PIP2-rich active zones, the sites of vesicle fusion where functionalities of Syt and other fusion machinery components are enhanced by PIP2 content (19, 34, 40). Driven by interactions between PIP2 and polybasic residues in the Syntaxin juxtamembrane LD, Syntaxin and PIP2 colocalize in ~70 nm clusters which may define sites of synaptic vesicle docking (65, 66). Our model predicts that vesicle docking mediated by *trans*-binding of the Syt ring occurs specifically at these PIP2- and Syntaxin-rich hot spots. Consistent with this picture, Syt promotes synaptic vesicle docking in Drosophila (18, 28) and chromaffin cells (31), and mutation of the Syt polylysine patch in mouse hippocampal neurons reportedly reduced the number of docked vesicles without affecting the total number of vesicles in the axon terminal (29).

### A model of Syt-mediated regulation of synchronous synaptic release

The present results are consistent with the proposed washer mechanism (43), and suggest that Syt rings may serve not only as membrane spacers but also as sensors that help direct synaptic vesicle docking to the active zone with high PIP2 content. In this picture, ATP-rich solution and negatively charged membranes compete for the Syt ring, a competition whose outcome depends on the PIP2 content of the membranes (Fig. 5*C*). (1) Prior to vesicle docking ATP-rich solution provides a more favorable environment than does the PIP2-free vesicle membrane, so that Syt C2AB domains unbind from the vesicle and organize into the halo that will later serve as landing gear and washer. (2) Once the vesicle approaches the PM, the PIP2-rich membrane provides a stronger pull for the Syt ring than does the ATP-rich solution, but only when PIP2 content exceeds a threshold. Thus, the Syt halo is spatially directed to dock at PIP2-rich sites that colocalize with other fusion components in the active zone. (3) Docking positions the Syt ring to clamp fusion as a spacer between vesicle and PMs, and likely by restraining the SNARE proteins via binding the primary interface identified in the Syt-SNARE complex crystal structure (23). The primary interface is positioned roughly opposite the polylysine patch, so it could bind the SNARE complex while simultaneously docking the vesicle to the PM (71). (4) When action potential-evoked Ca^2+^ influx disassembles the ring, the unrestrained SNAREs are free to fuse the membranes rapidly, facilitating neurotransmitter release through a fusion pore.

In this model, the Syt ring functions as a sensor that discriminates with high sensitivity between different levels of PIP2 and ATP. As a high sensitivity sensor, the Syt ring has a big advantage compared to individual Syt molecules, as rings have much greater binding energies. For example, compared to ATP-rich solution Syt rings prefer to bind PM with 20% local PIP2 content, but by a margin of only ~ kT per Syt molecule (i.e., trans-binding is preferred to the halo, Fig. 5*B*). Were unoligomerized Syt molecules the sensors, this advantage would be marginal, leaving a significant fraction of molecules unbound. By contrast, a 15 monomer Syt ring enjoys a substantial 15 kT binding advantage, so rings overwhelmingly prefer to bind a domain with 20% PIP2 content.

Thus, Syt rings allow regulation to be fine-tuned: the balance between halos and target membrane-bound rings is reversed by only a small change in conditions, such as local PIP2 content or ATP concentration. This translates to superior spatial regulation, ensuring that vesicles are directed to sufficiently PIP2-rich membrane domains with high fidelity.

## Materials and Methods

### Protein constructs, expression and purification

The Syt1 C2AB (residues 143 to 421) were expressed and purified using a pGEX6 vector with an N-terminal GST tag. Similar to previous studies, Escherichia coli BL21 gold (DE3) competent cells expressing Syt constructs were grown to an OD600 ~0.7-0.8, induced with 0.5 mM isopropyl β-D-1-thiogalactopyranoside (46, 72). The cells were harvested after 3.5 hours incubation at 37°C and suspended in lysis buffer containing 25 mM HEPES, pH 7.4, 400 mM KCl, 1 mM MgCl_2_, 0.5 mM TCEP, 4% Triton X-100, protease inhibitors. The samples were lysed, and the lysate was supplemented with 0.1% polyethylamine before being clarified by ultracentrifugation.

The supernatant was incubated with Glutathione-Sepharose beads overnight at 4°C. The beads were then washed with 20 mL of lysis buffer, followed by 20 mL of 25 mM HEPES, 400 mM KCl buffer containing with 2 mM ATP, 10 mM MgSO_4_, 1 mM DTT. Subsequently, the beads were re-suspended in 5 mL of lysis buffer supplemented with 10 μg/mL DNaseI, 10 μg/mL RNaseA, and 10 μL of benzonase (2000 units) and incubated at room temperature for 1 hour, followed by quick rinse with 10 mL of high salt buffer (25 mM HEPES, 1.1 M KCl, 1 mM DTT) to remove the nucleotide contamination. The beads were then washed with 20 mL of HEPES, 400 mM KCl buffer containing 0.5 mM EGTA to remove any trace calcium ions.

The N-terminal GST tag was cleaved by incubating with PreScission protease overnight at 4°C (46, 72). After eluting from the affinity beads, the protein was further purified by Mono-S anionic exchange (GE Healthcare, Marlborough, MA) chromatography.

### Isothermal Titration Calorimetry (ITC) measurements

ITC experiments were performed similarly to previously described.(73) Before the experiments, the protein Syt C2AB (residues 143 to 421) was purified by gel filtration using a Superdex 75 column (GE Healthcare Life Sciences), as before(73). Peak fractions were pooled and concentrated. Syt was then dialyzed overnight at 4°C. The concentration of dialyzed protein was determined using the Bradford assay with BSA as the standard. ATP was first dissolved in the dialysis buffer to 100 mM stock concentration and its pH was adjusted to 7.4, and filtered. Then ATP was diluted to ~800 μM for ITC experiments by adding the dialysis buffer.

ITC experiments were performed with a Microcal ITC200 instrument similarly to that was described before (74). Typically, about 200 μL of dialysis buffer or Syt solution was loaded into the sample cell, and about 60 μL ATP or PIP2 solution was loaded into the syringe. As described before(73, 74), ATP or PIP2 was first titrated in the dialysis buffer to make sure that no heat signal was generated, followed by titrations of ATP into Syt solution. The heat change from each injection was integrated, and then normalized by the moles of protein in the injection. Microcal Origin ITC200 software package was used to analyze the titration calorimetric data and obtain the stoichiometric number (*N*), the molar binding enthalpy (ΔH), and the association constant (*K_a_*). A nonlinear least-squares fit assuming a simple one site chemical reaction was used. The equilibrium dissociation constant (*K_D_*), the binding free energy (ΔG), and the binding entropy (ΔS) were calculated using the thermodynamic equations:

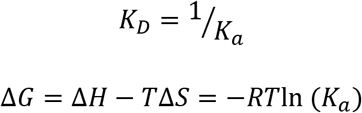

### Imaging of soluble Syt1 ring oligomers

Protein stock (50 μM) was diluted for 10 fold in MBS (20 mM MOPS, pH7.5, 15 mM KCl, 1 mM EGTA, 1 mM Mg(AC)2, 1 mM Mg.ATP, 1 mM DTT, 4% trehalose) at room temperature for 10 min. Diluted protein solution is further centrifuged at 10,000 x g for 10 min at 4°C to remove large aggregates and the supernatant (~8 μl) was applied to a continuous carbon-coated EM grid, which were glow-discharged for 10 s prior to application. After 1 min incubation, the grid was blotted dry with Whatman #1 filter paper, stained with 1% uranyl acetate and air-dried.

## Acknowledgements

We thank Shuyuan Wang and Rui Ma for valuable discussions. This work was supported by National Institute of Health grants GM071458 (J.E.R.) and R01GM117046 (B. O’S.), the HPC facilities operated by, and the staff of, the Yale Center for Research Computing, and Yale’s W. M. Keck Biotechnology Laboratory, as well as NIH grants RR19895 and RR029676-01 which helped fund the cluster.

## Author contributions

J.Z, J.E.R, B.O’S. designed the research; J.Z, F.L, Z.M. and B.O’S. performed the research and analyzed the data; Z.M., J.Z and B.O’S. wrote the paper.

