## Supporting Information for "Synaptotagmin rings as high sensitivity regulators of synaptic vesicle docking and fusion"

### Estimate of Syt-ATP, Syt-Syt, and Syt-membrane binding energy

We used a lattice model to calculate the Syt-ATP and Syt-Syt binding energies from our measured dissociation constant, and published values in the literature, respectively (1). The bulk solution in equilibrium was modeled on a lattice of site size $x$. In the dilute limit, the binding energy is

$$\epsilon = -kT \ln(x^3 k_d), \tag{S1.1}$$

where $k_d$ is the dissociation constant. Since the value of $x$ is determined by the smallest molecule in the reaction, it was chosen to be the size of an ATP molecule (~1 nm) for Syt-ATP binding and the size of a Syt molecule (~5 nm) for Syt-Syt binding.

For Syt-membrane binding, we used the same lattice model to calculate the binding energy from the molar partition coefficient measured by Kuo *et al*. (2). Lattice sites were divided into two groups, bulk and surface, to represent the aqueous and membrane phases. The ratio of the numbers of these sites depends on the area per lipid and the concentration of total lipids. The number densities of Syt in bulk ($\rho_b$) and on the membrane surface ($\rho_s$) satisfy $\rho_s = \rho_b \exp\left(\frac{\epsilon^{\text{syt-pm}}}{kT}\right)$. In the dilute limit, we have

$$\epsilon^{\text{syt-pm}} = kT \ln\left(\frac{K}{aA_0}\right) \approx 4.4\ \text{kT}. \tag{S1.2}$$

where $K = 160\ \text{M}^{-1}$ is the molar partition coefficient measured by Kuo *et al*. (2), $a = 5$ nm is the size of each site, and $A_0 = 0.64\ \text{nm}^2$ is the area per lipid (3).

The value of Syt-membrane binding energy at different salt concentration and lipid compositions was then extracted from electrostatic calculations. By approximating each polylysine patch to be a point charge $q$ and the lipid membrane to be an infinitely large surface with charge density $\sigma$, the electrostatic energy is

$$\epsilon^{\text{syt-pm}} = \frac{q\sigma}{4\pi\varepsilon_0\varepsilon_r}\int_0^\infty \frac{e^{-\frac{r}{\lambda}}}{r} 2\pi r\, dr = \frac{q\sigma\lambda}{2\varepsilon_0\varepsilon_r} \tag{S1.3}$$

where $\varepsilon_0$ and $\varepsilon_r$ are the electric permittivity and the relative permittivity of water, respectively (Table 1), $\lambda$ is the Debye length, and $r$ is the distance between the charge and a point on the membrane. Since $\lambda \propto [\text{salt}]^{-\frac{1}{2}}$, we have $\epsilon^{\text{syt-pm}} \propto \sigma[\text{salt}]^{-\frac{1}{2}}$.

### Model of Syt rings interacting with phospholipid membranes

**The Syt molecule.** The crystal structure of Syt (PDB ID: 2R83) shows that each C2 domain has an elongated shape with length $a = 5$ nm and width $b = 3$ nm (Fig. 1*B*). To approximate this geometry, we coarse-grained each C2 domain into two rigid spheres of radius $R_{\text{bead}} = \frac{b}{2} = 1.5$ nm with a fixed center-to-center separation of 2 nm. For simplicity, the C2A and C2B domains in each Syt are assumed closely packed and parallel to one another, i.e. the linker between the domains (residues 264-272) is effectively treated as a rigid connection. (In reality the linker has flexibility, relevant to the interaction between the C2A and C2B domains (4).)

In our model of the Syt molecule, we do not attempt to represent the highly charged juxtamembrane linker domain (LD) between the TMD and C2A (residues 80-140) (5). The LD could assist the binding of Syt rings to membranes since $\mathrm{Syt}^{\mathrm{CD}}$, the full cytosolic domain including the LD, oligomerizes into rings on monolayers more readily than does $\mathrm{Syt}^{\mathrm{C2AB}}$ (6). However, *in vivo* the LDs extend from the synaptic vesicle and presumably are not in close contact with the target membrane.

**Syt rings.** In Syt rings, each Syt subunit is assumed to bind two neighboring subunits at the two distal ends of the C2B domains, respectively (Fig. 1*C*). C2B domains are on the inside of the ring with their polylysine patches facing inward. The exact location of the polylysine patches with respect to the ring is unknown. EM reconstruction of Syt on monolayer tubes (Fig. 1*D*) suggests that polylysine patches are tilted by 30° from the mid-plane of the rings towards the membrane (Fig. 1*E*).

Since the twisting rigidity of Syt rings is unknown, for simplicity our model of the Syt ring neglects twisting motions that could rotate polylysine patches toward the membrane to which the ring is bound, so that contact between the polylysine patch and membrane could be achieved with less bending energy.

**Lipid membranes.** We used a force-based, triangular-mesh method based on (7) to model the lipid membranes. The profile of a membrane was represented by a triangular meshwork. Each node in the meshwork was connected to 4-12 neighboring nodes through tethered potentials, which is zero within a range of distance and infinity otherwise. Bond-flipping was included to capture membrane fluidity. In our simulations, planar membranes were represented by a $50$ nm $\times$ $50$ nm square meshwork with fixed boundaries. The meshwork contained 3600 membrane nodes, leading to an average bond length of $l_{\mathrm{mb}} \approx$ 0.9 nm and a tethering distance between 0.7 and 1.2 nm. Vesicles of 20 nm in radius were represented by a geodesic mesh generated from 6 subdivisions of an icosahedron, resulting 10242 nodes and an average bond length of $l_{\mathrm{ves}} \approx$ 0.8 nm. The tethering distance was between 0.6 and 1 nm.

The energy from bending and tension of the membrane is summed over all membrane nodes. The bending energy of the membrane is

$$F_{\mathrm{bend}}^{\mathrm{mb}} = \frac{\kappa}{2}\int (2H_i)^2 dA = \frac{\kappa}{2}\sum_i (2H_i)^2 A_i, \qquad \text{(S2.1)}$$

where $\kappa$ is the bending modulus of the membrane (Table 1), $H_i$ is the mean curvature of the $i^{\mathrm{th}}$ membrane node calculated following Ref. (8), and $A_i = \frac{1}{3}\sum_{j,k} A_{\Delta ijk}$ is the average area represented by the node. The tension energy (chemical potential) is

$$F_{\mathrm{tens}}^{\mathrm{mb}} = \gamma \int dA = \gamma \sum_i A_i, \qquad \text{(S2.2)}$$

where $\gamma$ is the membrane tension (Table 1).

**Syt ring-membrane interactions.** To calculate the electrostatic interactions between a Syt ring of $N$ subunits and a membrane, we treat each polylysine patch as a point charge $q$ and assume a uniform charge density $\sigma$ on the membrane. For our discrete membrane mesh, the amount of charges on the $i^{\mathrm{th}}$ node is $\sigma A_i$. From Eq. S1.3, the electrostatic energy between the $i^{\mathrm{th}}$ node of the membrane and the $j^{\mathrm{th}}$ Syt monomer is

$$F_{\mathrm{elec}}^{i,j} = \frac{q\sigma}{4\pi\varepsilon_0\varepsilon_r N}\frac{A_i}{r_{ij}} e^{-\frac{r_{ij}}{\lambda}} = \frac{\epsilon^{\text{syt-mb}}}{2\pi\lambda N}\frac{A_i}{r_{ij}} e^{-\frac{r_{ij}}{\lambda}}, \qquad \text{(S2.3)}$$

where $r_{ij}$ is the distance between the $i^{\text{th}}$ membrane node and the $j^{\text{th}}$ Syt monomer. A cutoff distance $r_{\text{cut}}$ is set for the electrostatic interaction so that $F_{\text{elec}}^{i,j}(r) = F_{\text{elec}}^{i,j}(r_{\text{cut}})$ for any $r < r_{\text{cut}}$. For a given membrane mesh, the value of $r_{\text{cut}}$ is calculated as follows: when a polylysine patch is $r_{\text{cut}}$ away from a node of an infinitely large flat membrane, the sum of electrostatic energies between the polylysine patch and all the mesh nodes should be $\epsilon^{\text{syt-mb}}$. By comparing the values of $\epsilon^{\text{syt-pm}}$ and $\epsilon^{\text{syt-ves}}$ to the energies calculated from our model, we find $r_{\text{cut}} = 0.7$ nm for Syt binding a PM and $r_{\text{cut}} = 0.2$ nm for Syt binding a synaptic vesicle.

The steric effect between the Syt ring and the membrane is assumed to be elastic when Syt is in contact with the membrane. The steric energy between the $i^{\text{th}}$ membrane node and the $j^{\text{th}}$ Syt monomer is

$$F_{\text{steric}}^{i,j} = k_{\text{steric}}\left(\Delta r_{ij}\right)^2, \tag{S2.4}$$

where $k_{\text{steric}} = 100$ pN/nm is an arbitrary spring constant and $\Delta r_{ij} > 0$ is the penetration of the $i^{\text{th}}$ node into the 4 beads of the $j^{\text{th}}$ Syt subunit.

For a Syt ring binding to a membrane, the change in total free energy is the sum of all relevant energies as shown in Eq. 2 in the main text.

**Carbon-supported monolayers: hydrophobic energy contribution.** For Syt rings on carbon-supported monolayers, there is an additional hydrophobic energy between the monolayer and carbon support. The hydrophobic energy of the $i^{\text{th}}$ monolayer node is (9, 10)

$$F_{h,i} = -E_h^0\left(e^{-\frac{z_i}{d_h}} - 1\right)A_i, \tag{S2.5}$$

where $E_h^0$ is an unknown constant, $z_i$ is the monolayer-carbon distance of the node, and $d_h \approx 2$ nm is the decay length of the hydrophobic interactions.

**Obtaining forces from the interaction energies.** The force between a Syt ring and a membrane is calculated as spatial derivatives of the electrostatic and repulsive energies: $-\nabla(F_{\text{elec}}^{\text{ring-mb}} + F_{\text{steric}}^{\text{ring-mb}})$. For membranes, additional normal stress resulted from the bending and tension energies on the $i^{\text{th}}$ node has been found (11) to be

$$p_i = 2\kappa_b\nabla^2 H_i - \kappa_b(2H_i + c_0)(2H_i - c_0)H_i - 2\gamma H_i, \tag{S2.6}$$

where $\nabla^2$ is the Laplace-Beltrami operator on the membrane surface and is evaluated following Ref. (12), $c_0$ is the spontaneous curvature of the membrane, and the contribution from Gaussian curvature is neglected. The force on the $i^{\text{th}}$ node from membrane bending and stretching is thus $p_i A_i \hat{n}_i$, where the local normal of the membrane $\hat{n}_i$ is evaluated from the area-weighted average normal of triangles surrounding node-$i$.

For vesicles of fixed volume, an isotropic internal pressure was included to conserve the volume as follows:

$$p_{\text{pres}} = -\frac{1}{\beta}\ln\left(\frac{V_{\text{ves}}}{V_{\text{ves}}^0}\right), \tag{S2.7}$$

where $\beta = 4.6 \times 10^{-10}$ Pa$^{-1}$ is the isothermal compressibility of water, $V_{\text{ves}}$ is the current volume of the vesicle, and $V_{\text{ves}}^0 = \frac{4}{3}\pi R_{\text{ves}}^3$ is the volume of the undeformed vesicle. To calculate $V_{\text{ves}}$, we divide the vesicle volume into tetrahedrons, each of which consists of the center of the vesicle and a triangular face in the vesicle mesh. $V_{\text{ves}}$ is thus the sum of the volume of all the tetrahedrons.

The total force on the $i^{\text{th}}$ node is thus

$$\vec{f}_i = -\sum_{j=1}^{N} \nabla_i \left(F_{\text{elec}}^{i,j} + F_{\text{steric}}^{i,j}\right) + \left(p_i + p_{\text{pres}}\right) A_i \hat{n}_i - \frac{dF_{h,i}}{dz_i} A_i \hat{e}_z, \tag{S2.8}$$

where $\hat{n}_i$ is the normal direction of the membrane at the $i^{\text{th}}$ node and $\hat{e}_z$ is the normal direction of the carbon surface. Note that we set $p_{\text{pres}} = 0$ for monolayers and planar membranes and $F_{h,i} = 0$ for bilayers. The force on the $j^{\text{th}}$ Syt monomer comes from the electrostatic and repulsion forces from the membrane:

$$\vec{f}_j = -\sum_i \nabla_j \left(F_{\text{elec}}^{i,j} + F_{\text{steric}}^{i,j}\right). \tag{S2.9}$$

**Evolving the Syt-membrane system with Langevin dynamics.** The system is evolved following Langevin equation at zero temperature. Because of symmetry, we treat each Syt ring as a rigid body undergoing only translational motion. The translational friction coefficient of the ring is the sum of that from all beads in each Syt subunit: $\zeta_{\text{ring}} = 4N\zeta_{\text{bead}}$, where $\zeta_{\text{bead}} = 6\pi\eta R_{\text{bead}}$ is the translational friction coefficient of each bead and $\eta$ is the viscosity of water. The displacement of the ring in time $\Delta t$ is

$$\Delta\vec{r}_{\text{ring}} = \frac{\Delta t}{\zeta_{\text{ring}}} \sum_{j=1}^{N} \vec{f}_j. \tag{S2.10}$$

The motion of membrane is calculated as follows. We treat each membrane node as a disc of area $A_i = \pi R_i^2$, the friction coefficient for the node moving along the normal direction is (13)

$$\zeta_{i,\perp} = 16\eta R_i. \tag{S2.11}$$

The two-dimensional (2D) in-plane translational friction coefficient of membrane node-$i$ is equal to that of a disc-like membrane patch with radius $R_i$ moving inside the membrane:

$$\zeta_{i,\parallel} \approx \frac{4\pi\eta_m}{\text{In}(\eta_m/\eta R_i)}, \tag{S2.12}$$

where $\eta_m$ is the 2D viscosity of the membrane (14). The instantaneous velocity due to the force is

$$\vec{v}_i = \frac{1}{\zeta_{i,\perp}} \vec{f}_i \cdot \hat{n}_i \hat{n}_i + \frac{1}{\zeta_{i,\parallel}} \vec{f}_i \cdot (1 - \hat{n}_i \hat{n}_i). \tag{S2.13}$$

The displacement of membrane node-$i$ in $\Delta t$ is

$$\Delta\vec{r}_i = \vec{v}_i \Delta t. \tag{S2.14}$$

The value of $\Delta t$ in our simulations is chosen to satisfy 1) change in energy for each particle/membrane node in each $\Delta t$ is less than 1 kT, 2) numerical stability of differential terms such as $\nabla^2 H$. In our simulations, the value of $\Delta t$ is mainly constrained by the latter and is estimated as follows. The motion of membrane nodes due to $\nabla^2 H$ term should lead to a displacement much smaller than the membrane mesh size $l_{\text{mb}}$. The highest curvature of membrane is estimated to occur when the membrane wraps around each C2 domain, the maximum curvature of which is $\sim 1/R_{\rm bead}$ and thus $\nabla^2 H \sim 1/R_{\rm bead} l_{\rm mb}^2$. The displacement of membrane nodes in each $\Delta t$ is $\Delta r_i \sim \frac{2\kappa_b (A_i \nabla^2 H)\Delta t}{\zeta_{i,\perp}} \sim \frac{2\kappa_b \Delta t}{\zeta_{i,\perp} R_{\rm bead}} \ll l_{\rm mb}$ . In the simulations we chose $\Delta t = 0.3 \frac{\zeta_{i,\perp} R_{\rm bead}}{2\kappa_b l_{\rm mb}} \sim 2$ ps.

The simulation was written in C programming language with OpenMP and was performed on high performance computer clusters. Typical run time for the system to reach equilibrium is about 2 days using 8 CPU cores.

**Calculation of the size distribution of Syt rings in solution**

Syt monomers can either be monomeric or belong to an oligomeric ring. The equilibrium volume fraction of rings with $N$ subunits is given by the Boltzmann distribution,

$$\phi_{\rm ring}(N) = \exp\left[-\frac{F_{\rm ring}(N)}{kT} + \frac{\mu N}{kT}\right], \tag{S3.1}$$

where $F_{\rm ring}(N) = -N\epsilon^{\text{syt-syt}} + E^{\rm ring}(N)$ is the free energy of an $N$-ring, $E^{\rm ring}(N)$is the bending energy of the ring, and $\mu$ is the chemical potential of Syt monomers of length $a$. For a ring of radius $R = Na/2\pi$, the bending energy is

$$E^{\rm ring} = \frac{l_p kT}{2}\left(\frac{1}{R} - \frac{1}{R_0}\right)^2 2\pi R, \tag{S3.2}$$

where $l_p$ is the persistence length and $R_0$ is the spontaneous radius of curvature of Syt oligomers. Rewriting this expression in terms of $N = 2\pi R/a$ and $N_0 = 2\pi R_0/a$, for small deviations $|N - N_0| \ll N_0$ from the minimum at $N = N_0$, we have

$$E^{\rm ring}(N) = \frac{2\pi^2 l_p kT}{a}\left(\frac{1}{N} - \frac{1}{N_0}\right)^2 N \approx gkT(N - N_0)^2, \tag{S3.3}$$

where $g = 2\pi^2 l_p / N_0^3 a$ and we expanded about $N_0$ to leading order. Then, Eq. S3.1 becomes

$$\phi_{\rm ring}(N) \approx \exp\left[-g(N - N_0)^2 + \frac{N}{kT}(\epsilon^{\text{syt-syt}} + \mu)\right]. \tag{S3.4}$$

The total concentration of rings of all sizes is thus

$$\Phi_{\rm ring}^{\rm tot} = \int_1^\infty \phi(N)dN \approx \int_{-\infty}^\infty \phi(N)dN = \sqrt{\frac{\pi}{g}} \exp\left[\frac{1}{4g}\left(\frac{\epsilon^{\text{syt-syt}}+\mu}{kT}\right)^2 + N_0 \frac{\epsilon^{\text{syt-syt}}+\mu}{kT}\right]. \tag{S3.5}$$

With rough estimates of $N_0 \sim 20$ and $l_p \sim 10^2$ nm, the above result shows that $\Phi_{\rm ring}$ is very small for $\mu < -\epsilon^{\text{syt-syt}}$, and very large for $\mu > -\epsilon^{\text{syt-syt}}$. Only near the critical value $\mu = -\epsilon^{\text{syt-syt}}$ can $\Phi_{\rm ring}^{\rm tot}$ have a value of order unity. This leads to a critical concentration of monomers $\phi_{\rm cmc} = \exp\left(-\frac{\epsilon^{\text{syt-syt}}}{kT}\right)$: for any total concentration of Syt molecules above $\phi_{\rm cmc}$, the relation $\mu \approx -\epsilon^{\text{syt-syt}}$ requires the free monomer concentration $\phi_f = \exp\left(\frac{\mu}{kT}\right)$ to be pinned to $\phi_{\rm cmc}$. This concentration is analogous to the critical micelle concentration in solutions of surfactant molecules. Thus, with $\mu = -\epsilon^{\text{syt-syt}}$, from Eq. S3.1 the size distribution of Syt rings is

$$P_{\mathrm{ring}}(R) \sim \exp\left(-\frac{E^{\mathrm{ring}}(R)}{kT}\right). \tag{S3.6}$$

**Supplemental Figures**

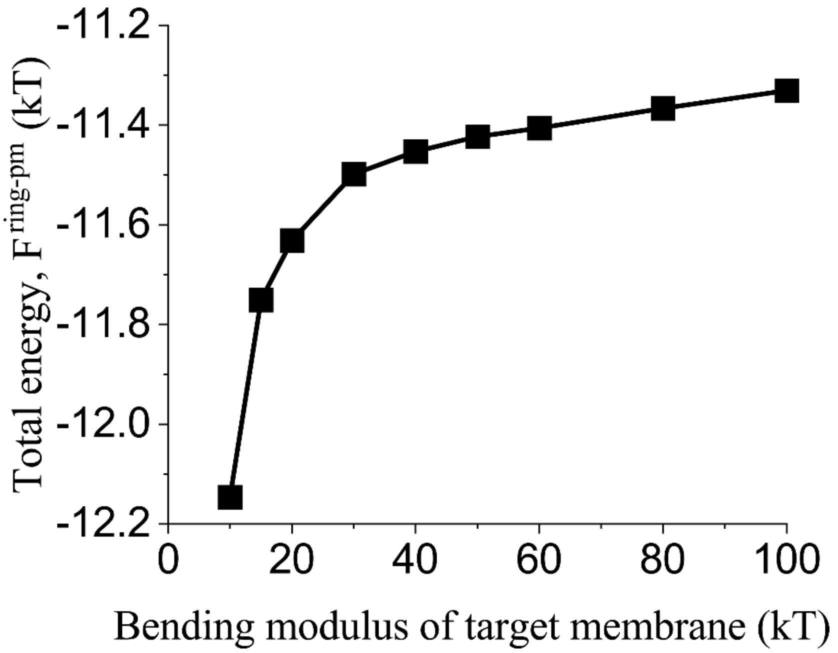


**Fig. S1.** Effect of bending modulus of membrane on the total free energy of a Syt ring of $N = 15$ binding to a planar membrane. The planar membrane has a lipid composition similar to that of target membrane (Table 2 in the main text).
